# On-ratio PDMS bonding for multilayer microfluidic device fabrication

**DOI:** 10.1101/450544

**Authors:** Andre Lai, Nicolas Altemose, Jonathan A. White, Aaron M. Streets

## Abstract

Integrated elastomeric valves, also referred to as Quake valves, enable precise control and manipulation of fluid within microfluidic devices. Fabrication of such valves requires bonding of multiple layers of the silicone polymer polydimethylsiloxane (PDMS). The conventional method for PDMS-PDMS bonding is to use varied base to crosslinking agent ratios between layers, typically 20:1 and 5:1. This bonding technique, known as “off-ratio bonding,” provides strong, effective PDMS-PDMS bonding for multi-layer soft-lithography, but it can yield adverse PDMS material properties and can be wasteful of PDMS. Here we demonstrate the effectiveness of on-ratio PDMS bonding for multilayer soft lithography. We show the efficacy of this technique among common variants of PDMS: Sylgard 184, RTV 615, and Sylgard 182.

## 1. Introduction

The integrated elastomeric valve is the transistor of microfluidic circuits. Its development has enabled a suite of fundamental microfluidic components, including logic gates [1], peristaltic pumps [1], active cell traps [2], microfluidic formulators [3], and button membranes for mechanical trapping of molecular interactions [4]. These components are used to construct integrated microfluidic devices, which have proven to be powerful research tools in quantitative biology. Applications of such devices include PCR [5], digital PCR [6], qPCR on chip [7], microfluidic bioreactors [8], high-throughput parallel analysis [9], protein crystallography [10] and single cell analysis [11,12]. A vital component of integrated elastomeric valve fabrication is multilayer soft lithography [13], a technique which involves bonding casted layers of polydimethylsiloxane (PDMS) which contain control and flow channels. While many methodologies exist to bond multiple layers of PDMS, including oxygen plasma [14], adhesive [15,16], and corona discharge [17], the standard method for PDMS-PDMS bonding is termed “off-ratio bonding”. This technique takes advantage of the fact that PDMS is a two-component elastomer consisting of a base, also known as potting compound, and a crosslinking agent. To perform off-ratio bonding, two layers of partially cured PDMS with varying base-to-crosslinker ratios are brought into contact (Fig. 1). Because each layer has an excess of one component (when compared to standard 10:1 ratios), a concentration gradient is created. This gradient is thought to drive the diffusive transport of reactive molecules across the bonding interface to irreversibly bond the two layers [13,18]. The primary advantage of off-ratio bonding over the aforementioned methodologies is that the bond is not permanent until thermal curing. This enables iterative adjustment of the layers to align control and flow channels on the two respective layers. With plasma and corona discharge, bonding occurs instantaneously when the layers are brought into contact. If the layers are misaligned, further manipulation to realign the layers cannot be performed. With adhesive bonding, repeated manipulation of the layers may unevenly distribute the adhesive or may inadvertently push adhesive into channels.

**Fig. 1.**
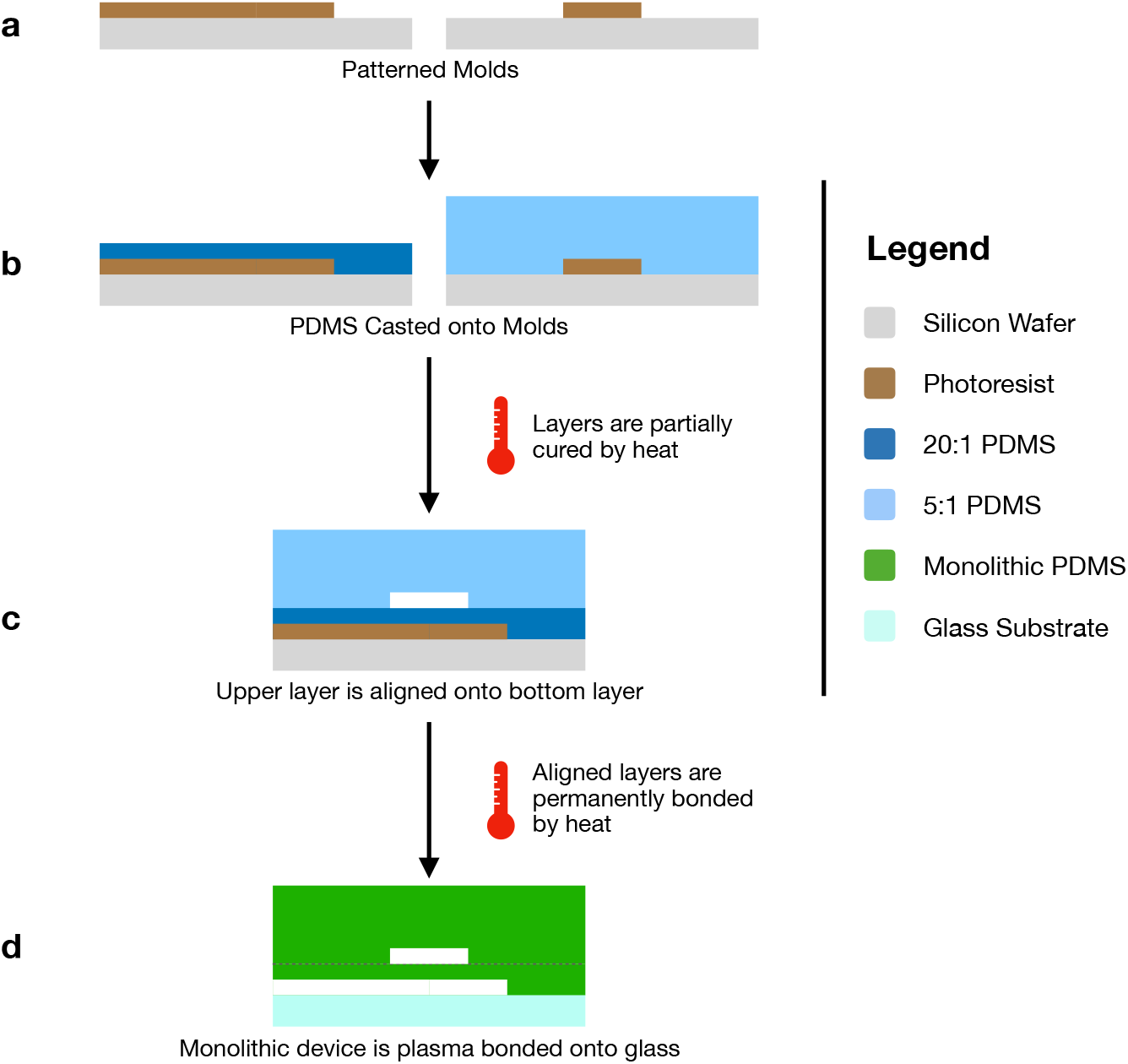
Multilayer soft-lithography using conventional off-ratio bonding: (a) Molds are patterned using photolithography; (b) 20:1 base-to-crosslinker PDMS is spin-coated onto the flow layer mold (left); 5:1 base-to-crosslinker PDMS is poured onto the control layer mold (right); (c) Each layer is partially thermally cured; the top layer is then aligned onto the bottom layer; (d) Permanent bonding between the aligned layers is achieved by a longer thermal curing step. During this process, a concentration gradient is thought to drive the diffusive movement of molecules across the bonding interface to complete the bonding process. The device is then plasma bonded onto a glass substrate.

Off-ratio bonding provides a seamless bond between the control and flow layers. This technique can hold up to 72 PSI (500 kPa) [19], well above a normal microfluidic operating range of 20 - 45 PSI (138 - 310 kPa). However, there are some drawbacks to using non-standard ratios of base to crosslinking agent. First, the two components of PDMS are typically manufactured and distributed for use with a 10:1 ratio of base to crosslinking agent. Because the thicker of the two layers must have the lower ratio, typically 5:1, excess base is accumulated and ultimately wasted. Second, the material properties of 5:1 and 20:1 PDMS differ from the properties of 10:1 PDMS as specified by the manufacturers. For many applications, this may not be a substantial issue; physical material characteristics for 5:1 and 20:1 PDMS have been reported [20,21]. However, when layers of 5:1 and 20:1 PDMS are brought into contact for off-ratio bonding, the resultant material properties of the monolithic PDMS are poorly specified because diffusion of excess crosslinking agent across the bonding interface alters the base-to-crosslinker ratio of the final PDMS [22]. We have also observed differing optical properties when using non-standard ratios of PDMS (see Supplemental Figure S1). A final significant issue with off-ratio bonding is the potential toxicity of excess PDMS crosslinking agent with certain cell types. PDMS with base-to-crosslinker ratios less than the conventional 10:1 ratio can cause certain cell types to detach and may inhibit them from reaching confluence [23].

Here we present an alternative PDMS-PDMS bonding technique that helps mitigate the aforementioned issues with off-ratio bonding. We demonstrate that a stable and effective bond can be produced using a standard “on-ratio” mixture of 10:1 base to crosslinking agent. We use this technique with three common types of PDMS: Sylgard 184, RTV 615, and Sylgard 182 and compare the bond strength to off-ratio bonding using Sylgard 184 and RTV 615. When utilized to create microfluidic devices with integrated elastomeric valves, all five techniques perform equally well within a normal microfluidic operating range of 20 - 45 PSI (138 - 310 kPa). Likewise, there were minimal significant differences in valve responsiveness between the devices fabricated with off-ratio bonding and on-ratio bonding. On-ratio bonding is a robust protocol that permits repeated manipulation of layers during alignment while eliminating disadvantages such as excess PDMS base, equivocal material properties, and potentially limited biocompatibility.

## 2. Methods

### 2.1 Mold Fabrication

Molds were created using standard photolithography. In total, two control layer molds and one flow layer mold were fabricated. Photomasks were designed using AutoCAD and commercially produced using a 25400 dpi printer (CAD/Art Services, Inc., Bandon, Oregon). For the control layer molds, a dummy layer of SU8-2005 (MicroChem Corp., Westborough, MA) was spin-coated at 3500 rpm onto 10 cm silicon wafers (University Wafer, Boston, MA) and subsequently cured to promote adhesionof the mold features. SU8-2025 Photoresist (MicroChem Corp., Westborough, MA) was then spin-coated (Brewer Science, Rolla, MO) on top of the dummy layer at 2500 rpm for a feature height of 25 μm. Exposure was performed using a UV Aligner (OAI, San Jose, CA). Specific baking temperatures, baking times, exposure dosages, and development times followed the MicroChem data sheet. For the flow layer mold, AZ 40XT (Integrated Micro Materials, Argyle, TX) positive photoresist was spin coated at 3000 rpm for a feature height of 20 μm. Specific baking temperatures, baking times, and development times followed a previously published protocol [12]. To round the features, the wafer was baked on a hot plate for 1 min at 65°C, for 1 min at 95°C, and for 1 min 135°C to reflow the photoresist.

### 2.2 Device Fabrication

Soft-lithography procedures [24] were employed to fabricate each device. In order to test the bond interface, a thick control layer with microfluidic features was bonded to a thin dummy layer without microfluidic features (Fig. 2). In total, five device variants were fabricated: three device variants employed on-ratio bonding (our method) and two device variants employed standard off-ratio bonding. The PDMS composition of each layer in each device variant is detailed in Table 1. For each experiment, a total of 50 devices were fabricated (10 replicates per variant). Control molds and blank wafers (for the dummy layer) were first treated with trichloromethlysilane (Sigma-Aldrich, St. Louis, MO) in a vacuum chamber for 20 min. Each PDMS variant was then mixed for 2 min with a vertical mixer and then degassed in a vacuum chamber. Sylgard variants were degassed for 45 min and RTV615 variants were degassed for 90 min. PDMS was then poured onto the control molds and degassed for an additional 5 min. PDMS was spin-coated (Specialty Coating Systems, Indianapolis, IN) onto the blank wafers. A two-step spin protocol was utilized. In the first step, PDMS was spun at 500 rpm, using an acceleration of 100 rpm/s and a 2 sec dwell time. In the second step, PDMS was spun to a final speed dependent on the PDMS variant and detailed in Table 1, using an acceleration of 500 rpm/s and a 60 sec dwell time. The second step spin speed was optimized for a thickness of 55 μm (see Supplemental Figure S2). The layers were then baked at 70°C using a forced air convection oven (Thermo Fisher Scientific, Waltham, MA) to partially cure the PDMS. Specific baking times for each layer are detailed in Table 1 (see Supplemental Section S1 and Section S2). After baking, the control layers were cut out, hole punched (SCHMIDT Technology, Cranberry Township, PA) (see Supplemental Section S3), and aligned onto the dummy layer. A final post-alignment baking step was performed at 70°C for 120 min to complete the bonding. The bonded devices were then mounted onto glass coverslips (VWR International, LLC, Radnor, PA) using oxygen plasma (Plasma Equipment Technical Services, Brentwood, CA), with five devices of the same variant on each coverslip. Other device designs, used to test pressure with large chamber geometries or to test valve responsivity, were fabricated according to the same protocol, with different master molds.

**Fig. 2.**
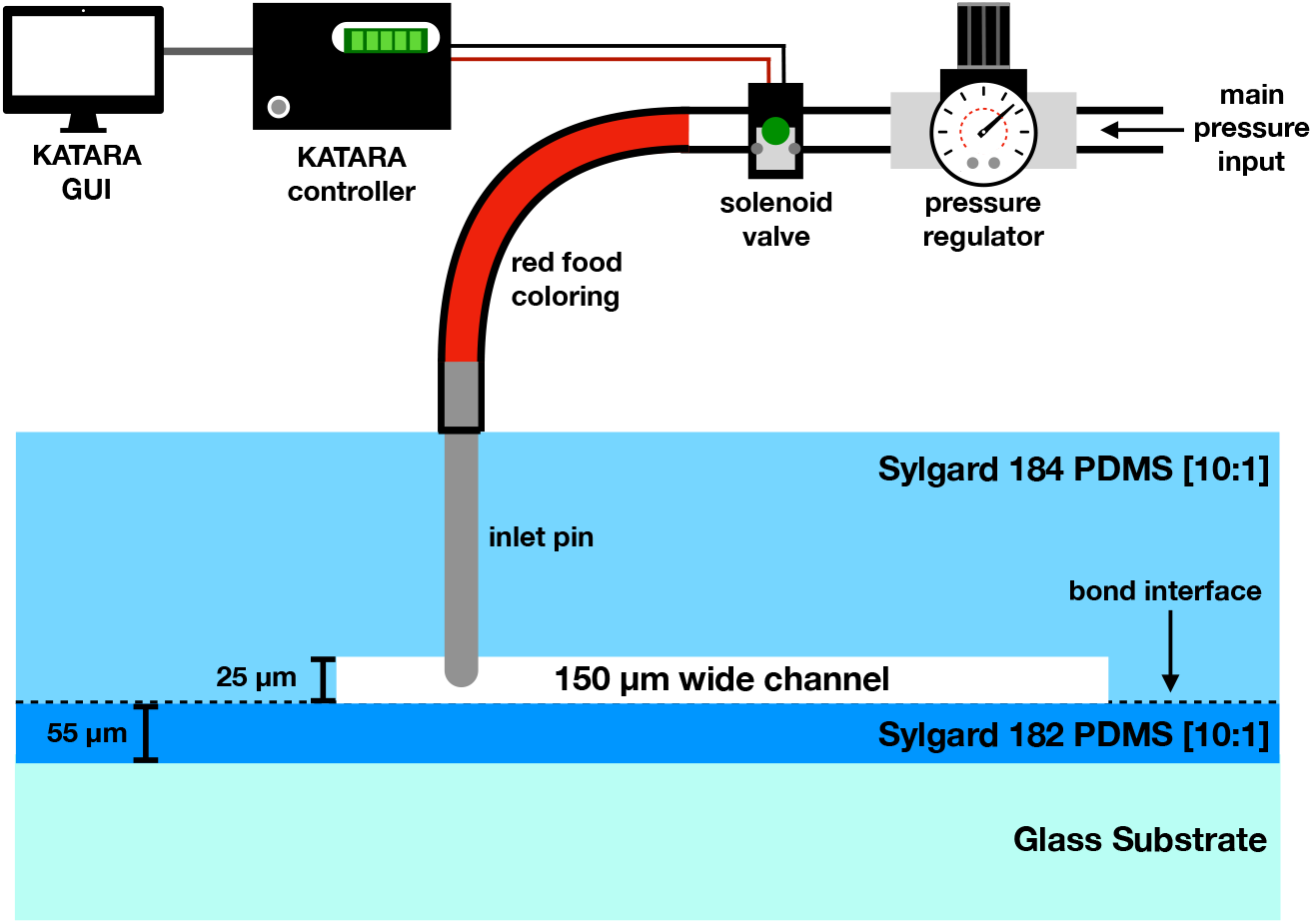
Schematic of experimental set-up: pressure from main pressure lines are controlled by a pressure regulator and actuated using a solenoid valve connected to the KATARA microcontroller and software GUI. Using this pressure, each device was filled with red dye to test the bond strength.

**Table 1.** PDMS Specifications, Baking Times, and Thin Layer Spin Speeds of Each Variant:

| Variant # | Variant Composition |  | Baking Time at 70°C (min) | Spin Speed (rpm) |
| --- | --- | --- | --- | --- |
| 1 | Thick Layer | Sylgard 184 10:1 | 25 | n/a |
|  | Thin Layer | Sylgard 182 10:1 | 32 | 1400 |
| 2 | Thick Layer | Sylgard 184 10:1 | 25 | n/a |
|  | Thin Layer | Sylgard 184 10:1 | 14 | 1400 |
| 3 | Thick Layer | Sylgard 184 5:1 | 19 | n/a |
|  | Thin Layer | Sylgard 184 20:1 | 19 | 1900 |
| 4 | Thick Layer | RTV 615 10:1 | 19 | n/a |
|  | Thin Layer | RTV 615 10:1 | 10 | 2000 |
| 5 | Thick Layer | RTV 615 5:1 | 16 | n/a |
|  | Thin Layer | RTV 615 20:1 | 16 | 2000 |

### 2.3 Experimental Set-Up

The experiment was conducted blind: prior to connecting the pressure inlets to the devices, a neutral colleague unaffiliated with the experiment assigned each device an alphabetical code corresponding to which bonding variant was used. This ensured that unintentional bias could not be imparted when recording bond delamination. A control channel in each device was filled with red dye (Fisher Science Education, Nazareth, PA) and pressurized to 20 PSI (138 kPa) (Fig. 2). This was done using an external control set-up that consisted of a pressure regulator connected to the main pressure line. This regulated pressure was connected to a solenoid valve array (Pneumadyne, Plymouth, MN) to individually address each valve with a control pressure. The solenoid valve array was controlled by the KATARA microfluidics controller and software [25]. See Supplemental Figure S3 for more information about the experimental set-up.

### 2.4 Device Pressure Testing

All variants were tested in parallel (Fig. 3). For each device, one microfluidic channel measuring 150 μm in width was pressurized at steady pressure for 30 min. After 30 min of sustained pressure, pressure was repeatedly actuated at 0.5 Hz for another 30 min to simulate valves opening and closing. This one-hour test was performed at 30 PSI (207 kPa), 45 PSI (310 kPa), and 60 PSI (414 kPa).

**Fig. 3.**
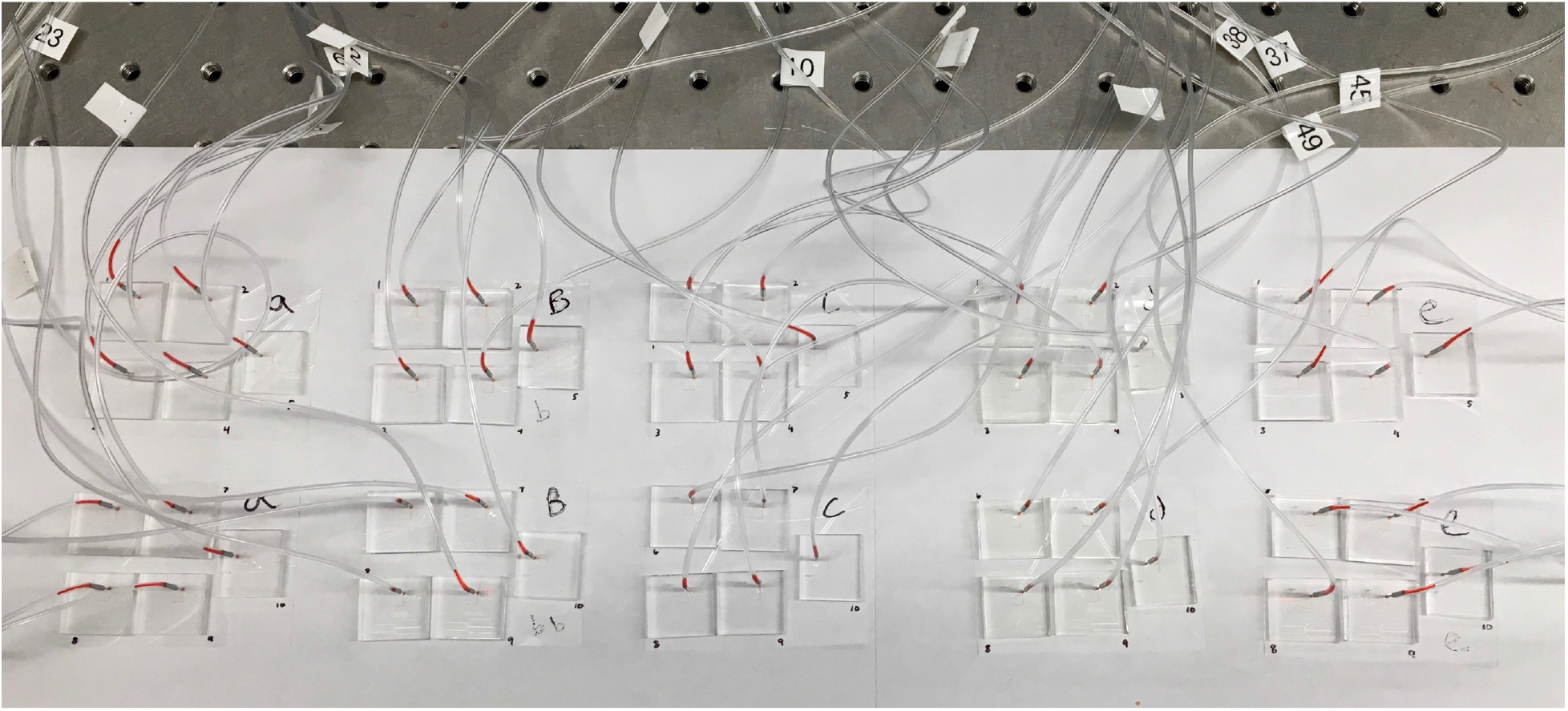
Image of All 50 Devices Being Tested in Parallel: five variants, with ten replicates per variant, were connected to 50 external solenoid valves. These external solenoid valves were connected to a common input pressure line controlled by a pressure regulator.

Qualitative examination of the red dye clearly showed whether or not delamination between the flow and control layers occurred, which is indicative of bond failure. When the device delaminates, red dye would escape between the two layers. After delamination, red dye would no longer be present in the clear tubing or in the microfluidic channels, and a path of delamination could be traced from the microfluidic channel to the device perimeter. In some cases, the device failed by another mechanism not indicative of bond failure. These failure modes include splitting of the bulk PDMS or ejection of the inlet pin from the device itself.

Subsequent experiments to further validate the bond strength of the devices that used on-ratio bonding involved implementing 7 mm diameter chamber geometries (Fig. 4). Previous studies have used large chambers to test bond strength in order to make the bond more susceptible to delamination [19]. With these large chambers, we could effectively test the bond strength of the devices to the point of failure without the need for pressure sources that could output higher pressures. These devices were tested using the same methodology described for the normal geometry devices, but without the 30 min of on/off actuation.

**Fig. 4.**
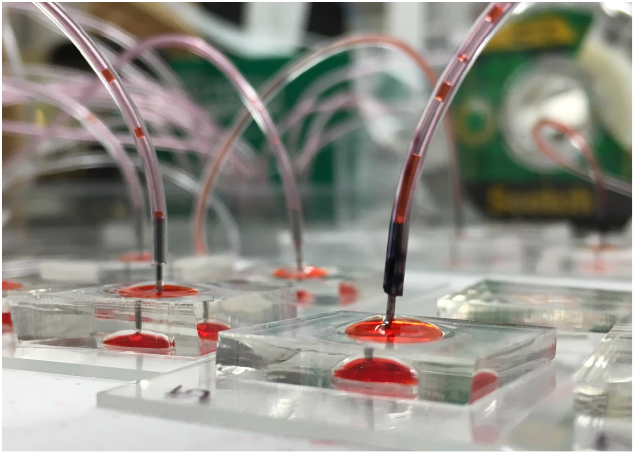
Images of Device with 7 mm Circular Geometry: devices with the 7 mm circular chamber geometry were pressurized with red dye.

### 2.5 Valve Response Testing

The valves of devices fabricated using 182/184 on-ratio bonding protocol were compared to valves of devices fabricated using the standard 184 off-ratio protocol. These devices were fabricated using the same set of molds. The PDMS thicknesses of the flow layers of each fabrication type were controlled to a height of 55 μm; this ensured that membrane thickness would not be a variable in valve responsivity. To test the opening and closing times of the valves, green dye (Fisher Science Education, Nazareth, PA) was filled into a single flow channel, while water was dead-end filled into an overlapping control channel. The device was placed under a scanning confocal microscope (Olympus, Center Valley, PA). The valve was opened and closed under 20 PSI (138 kPa) of pressure repeatedly at 4 Hz. As the valve closes, green dye in the underlying channel is displaced (Fig. 5). When the valve returns to the open position, the green dye in the underlying channel is restored. A transmission detector recorded the series of valve responses. The transmission intensity time series was then processed in R (R Foundation). Response times were found by determining the time required for the signal to transition between 10% and 90% of the max-min intensity.

**Fig. 5.**
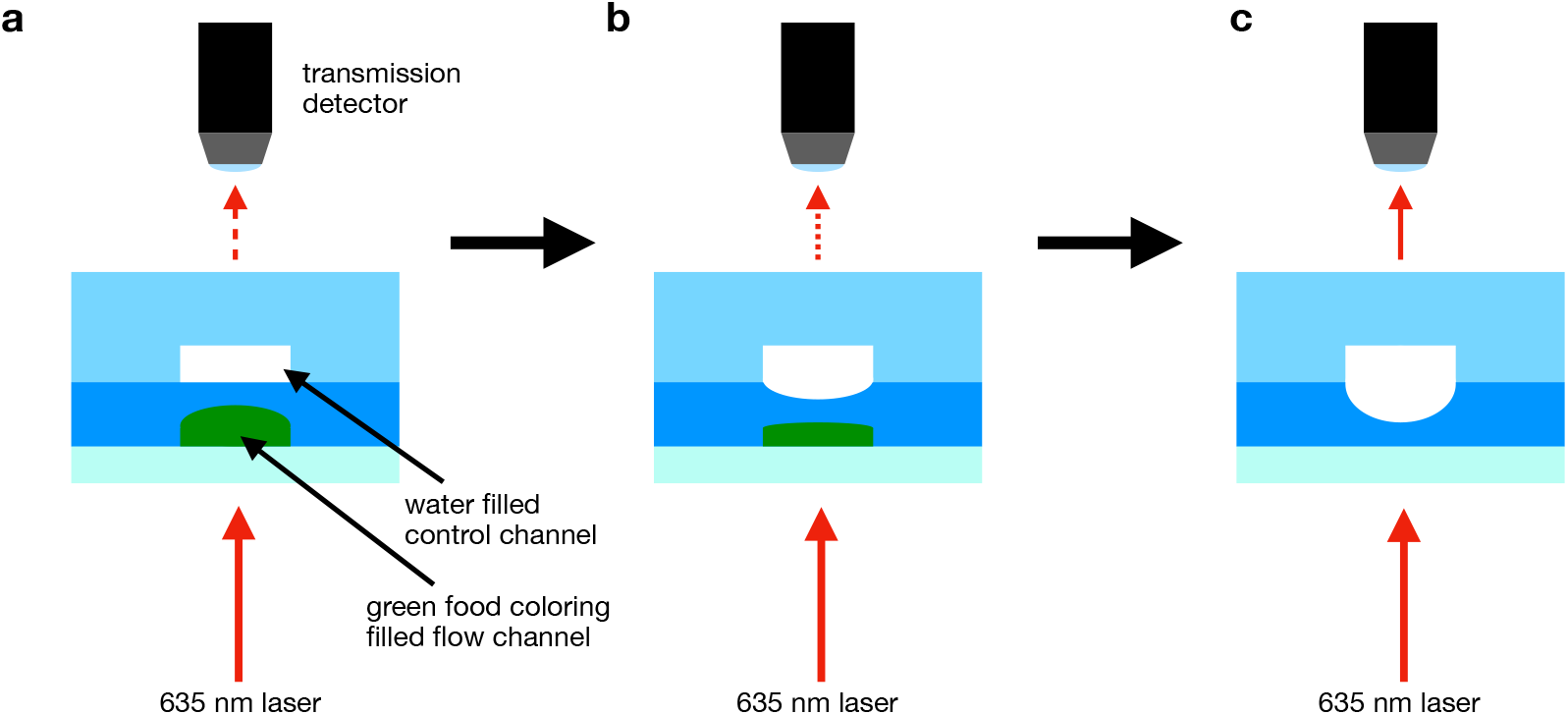
Schematic of Valve Responsivity Measurement: (a) The valve is fully open. Green dye fills the lower flow channel, and water fills the upper control channel. Transmission is at a minimum; (b) The upper control channel is pressurized, and the valve is in the process of closing. The green dye is being displaced. Transmission is increasing; (c) The valve has fully closed the flow channel. All green dye has been displaced and transmission is at a maximum.

## 3. Results and Discussion

### 3.1 Bond Performance

In total, 20 devices for each variant were tested. Pressurizing single microfluidic channels with normal geometries (150 μm width and 25 μm height) resulted in no delamination at 30 and 45 PSI. At 60 PSI, five bond failures for 184 on-ratio devices and one bond failure for RTV on-ratio devices occurred (Fig. 6). A bond failure is defined as a visible delamination of the two layers. This means that a path of delamination can be traced from the channel to the perimeter of the device. A “failure not due to bonding” is any other failure that rendered the device unsuitable for pressure testing e.g., plasma bonding error, inlet pin ejection, bulk PDMS splitting, etc.

**Fig. 6.**
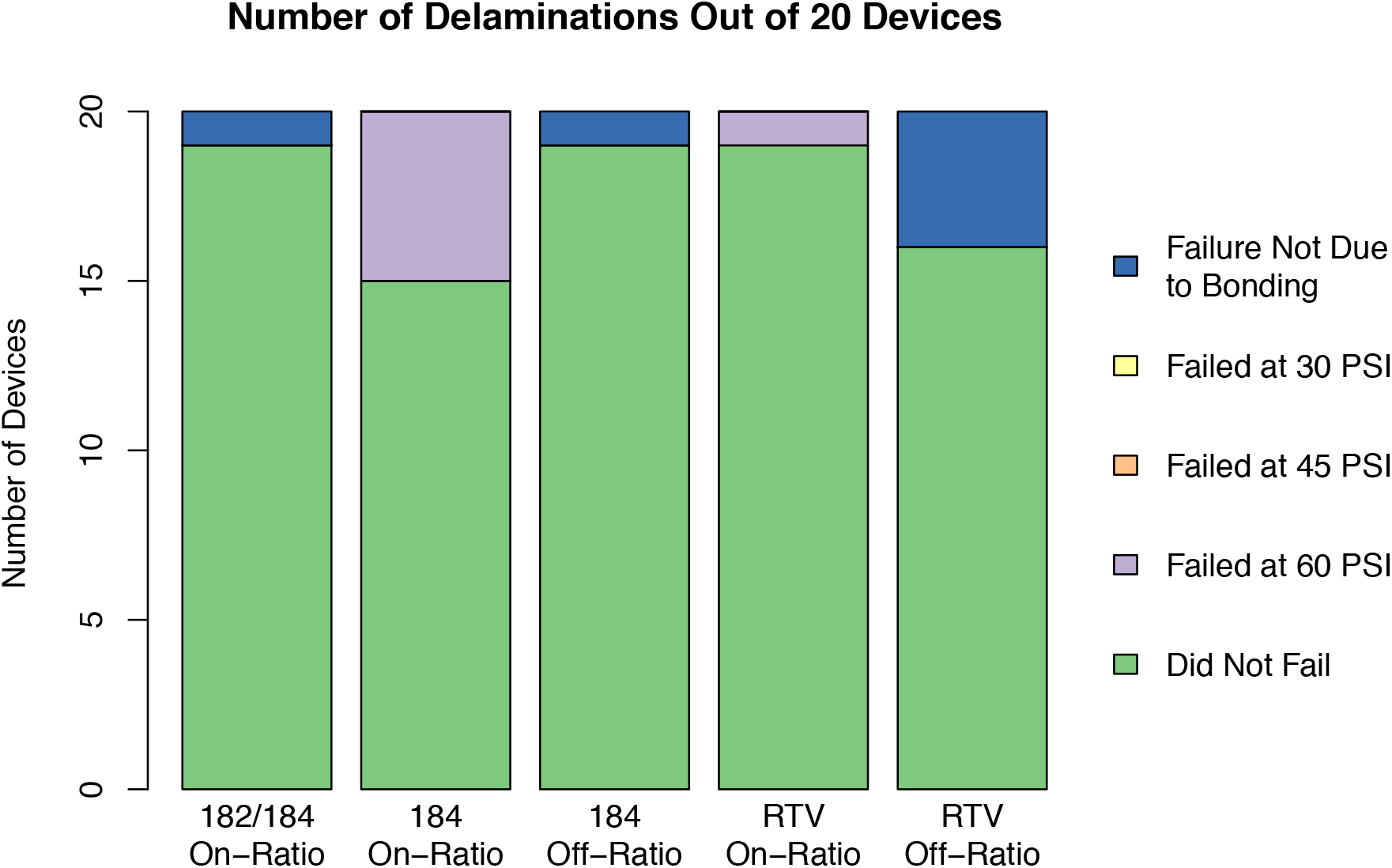
Results of Bond Performance. Pressurizing channels 150 μm wide and 25 μm tall resulted in no delamination at 30 and 45 PSI. At 60 PSI, 5/20 and 1/20 devices delaminated for 184 on-ratio and RTV on-ratio, respectively.

The majority of devices fabricated using the on-ratio protocol were able to withstand up to 60 PSI. In order for such a strong bond to exist, covalent interactions must exist across the bonding interface. Secondary forces between polymer chains are not sufficient for creating a strong, permanent bond [26]. To have covalent interactions across the bonding interface, polymer chains would have to bridge the bonding interface. Consequently, our results support the theory that there is sufficient self-diffusion of polymer chains between two layers of PDMS of identical composition to create an autohesive bond that can withstand normal microfluidic operating pressures of up to 60 PSI. While off-ratio bonding may provide an additional driving force for diffusion of polymer molecules across the bonding interface, our results demonstrate that this driving force may not be necessary to create a bond sufficient for multilayer soft lithography.

### 3.2 Burst Pressure Testing

To further test the limits of our bonding technique, pressure tests were performed on two variants: 182/184 on-ratio and 184 on-ratio, using large 7 mm circular diameter chambers. Despite the larger chambers, the majority of chips were able to withstand up to 45 PSI (Fig. 7). Standard microfluidic operating pressure is within the range of 20 - 45 PSI. This result demonstrates that even with extreme geometries, on-ratio bonding is effective for most microfluidic applications.

**Fig. 7.**
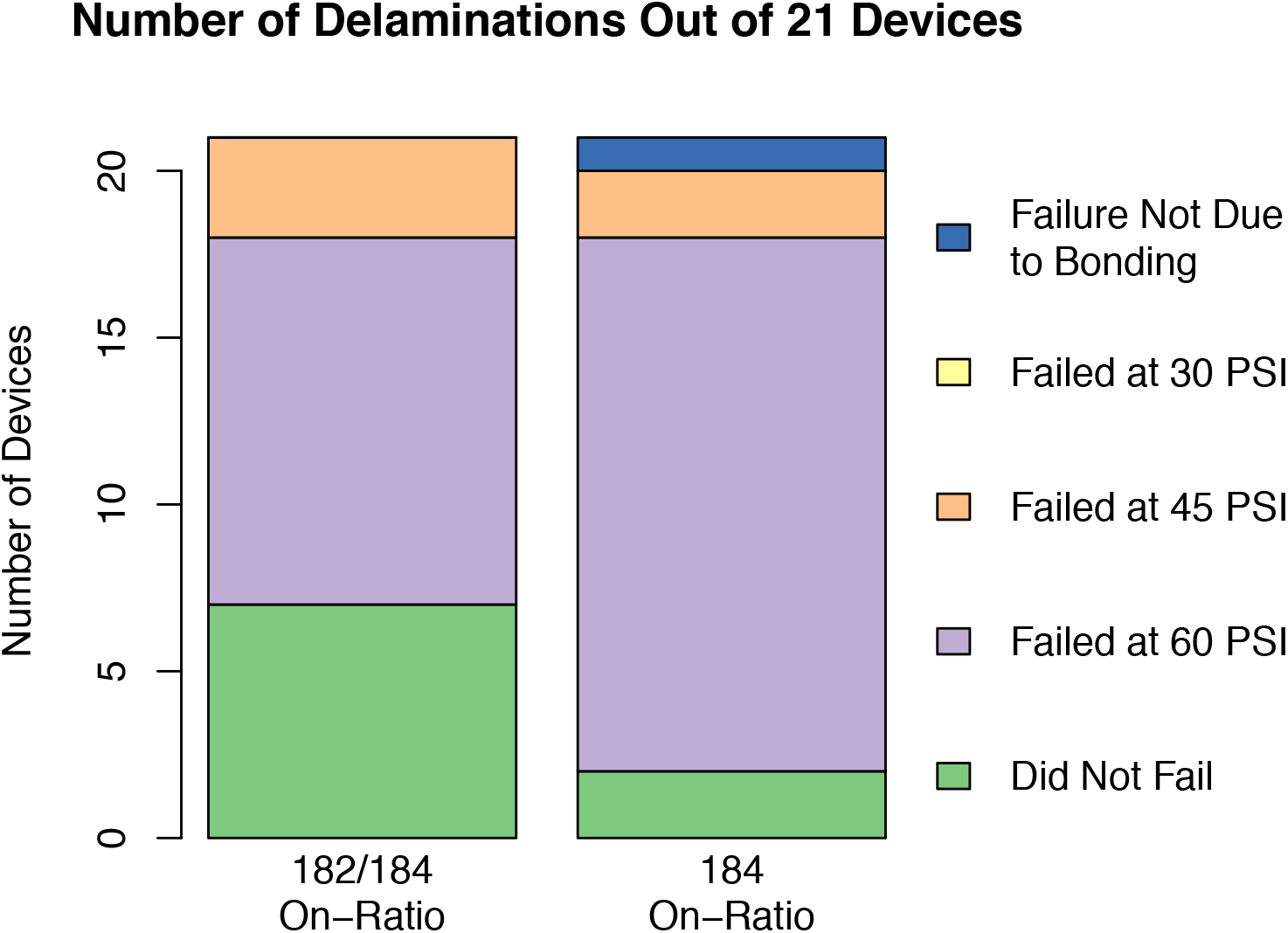
Large Geometry Bond Performance: Chambers with a diameter of 7 mm were used to further verify the strength of on-ratio bonding. For both variants tested (182/184 on-ratio and 184 on-ratio), the majority of devices withstood up to 45 PSI.

### 3.3 Valve Responsivity

The valve responsiveness of the 182/184 on-ratio devices had a closing time comparable to the 184 off-ratio devices, both averaging about 10 ms. However, for opening times, the 184 off-ratio devices performed over twice as fast, averaging about 2.6 ms versus 6.6 ms for the 182/184 on-ratio devices (Fig. 8). These differences are on the order of a few milliseconds and have negligible effect on the real-world operation of the devices.

**Fig. 8.**
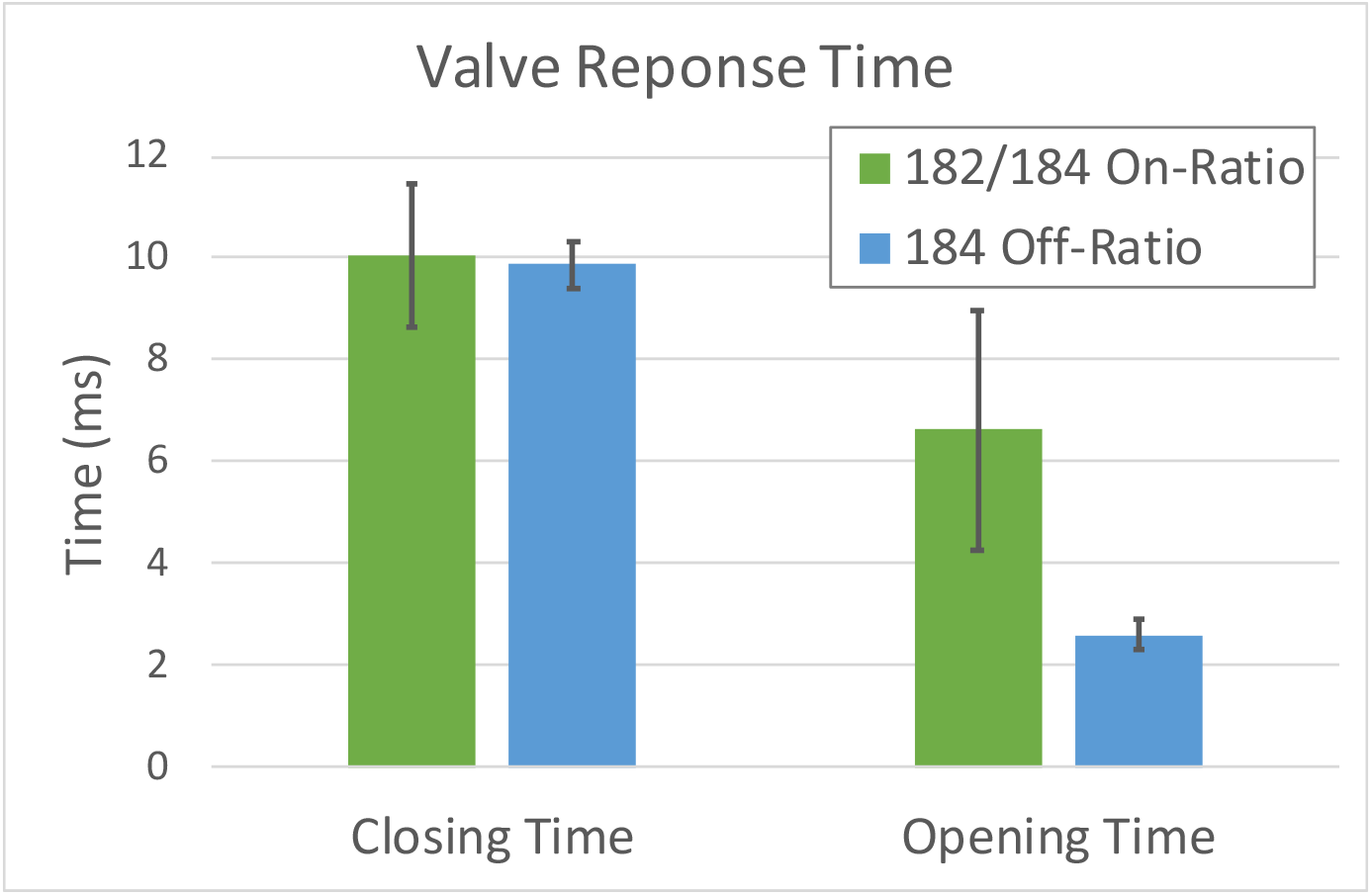
Comparison of Valve Opening and Closing Time: The 182/184 on-ratio devices had a closing time similar to that of the 184 off-ratio devices, with averages at 10.0 ms and 9.9 ms for the on-ratio devices and off-ratio devices, respectively. For opening times, the 184 off-ratio devices were over twice as fast than the 182/184 on-ratio devices, averaging 2.6 ms versus 6.6 ms for the 182/184 on-ratio devices. Error bars represent 2 standard errors.

## 4. Conclusion

The use of on-ratio bonding provides an effective alternative to off-ratio bonding at normal operating pressure range of 20 - 45 PSI (138 - 310 kPa). This technique is robust and compatible with fabricating integrated elastomeric valves. Valves fabricated using this novel technique were shown to be comparable in closing times to valves fabricated using off-ratio techniques. Overall, on-ratio bonding eliminates the problem of excess PDMS base, helps maintain normal material characteristics, and is potentially more biocompatible than devices fabricated using off-ratio techniques [23]. Furthermore, on-ratio bonding permits repeated manipulation of the layers during alignment before permanent bonding, a critical component of valve fabrication.

## Acknowledgements

We would like to thank Gabriel Dorlhiac for his assistance with measurements involving the scanning confocal microscope and spectrophotometer. We would also like to thank Paul Lum whose discussion inspired this investigation. Finally, we would like to thank all members of the Streets Lab for their support. Research reported in this publication was supported by the University of California, Berkeley, Department of Bioengineering and the National Institute of General Medical Sciences of the National Institutes of Health [Grant Number R35GM124916]. Andre Lai is supported by the UC Berkeley Haas Scholars Program. Nicolas Altemose is a Howard Hughes Medical Institute Gilliam Fellow. Aaron Streets is a Chan Zuckerberg Investigator.

## References

[1] T. Thorsen, S.J. Maerkl, S.R. Quake, Microfluidic large-scale integration, Science. 298 (2002) 580–584. doi:10.1126/science.1076996.

[2] Y. Marcy, C. Ouverney, E.M. Bik, T. Losekann, N. Ivanova, H.G. Martin, E. Szeto, D. Platt, P. Hugenholtz, D.A. Relman, S.R. Quake, Dissecting biological “dark matter” with single-cell genetic analysis of rare and uncultivated TM7 microbes from the human mouth, Proc. Natl. Acad. Sci. 104 (2007) 11889–11894. doi:10.1073/pnas.0704662104.

[3] C.L. Hansen, M.O.A. Sommer, S.R. Quake, Systematic investigation of protein phase behavior with a microfluidic formulator, Proc. Natl. Acad. Sci. 101 (2004) 14431–14436. doi:10.1073/pnas.0405847101.

[4] S. Rockel, M. Geertz, S.J. Maerkl, MITOMI: A microfluidic platform for in vitro characterization of transcription factor-DNA interaction, Methods Mol. Biol. 786 (2012) 97–114. doi:10.1007/978-1-61779-292-2_6.

[5] J. Liu, M. Enzelberger, S. Quake, A nanoliter rotary device for polymerase chain reaction, Electrophoresis. 23 (2002) 1531. doi:10.1002/1522-2683(200205)23:10<1531::AID-ELPS1531>3.0.CO;2-D.

[6] E.A. yOttesen, W.H. Jong, S.R. Quake, J.R. Leadbetter, Microfluidic digital PCR enables multigene analysis of individual environmental bacteria, Science. 314 (2006) 1464–1467. doi:10.1126/science.1131370.

[7] A.K. White, M. VanInsberghe, O.I. Petriv, M. Hamidi, D. Sikorski, M.A. Marra, J. Piret, S. Aparicio, C.L. Hansen, High-throughput microfluidic single-cell RT-qPCR, Proc. Natl. Acad. Sci. 108 (2011) 13999–14004. doi:10.1073/pnas.1019446108.

[8] A.J. Kaestli, M. Junkin, S. Tay, Integrated platform for cell culture and dynamic quantification of cell secretion, Lab Chip. 17 (2017) 4124–4133. doi:10.1039/c7lc00839b.

[9] S. Kim, J. De Jonghe, A.B. Kulesa, D. Feldman, T. Vatanen, R.P. Bhattacharyya, B. Berdy, J. Gomez, J. Nolan, S. Epstein, P.C. Blainey, High-throughput automated microfluidic sample preparation for accurate microbial genomics, Nat. Commun. 8 (2017) 13919. doi:10.1038/ncomms13919.

[10] C.L. Hansen, E. Skordalakes, J.M. Berger, S.R. Quake, A robust and scalable microfluidic metering method that allows protein crystal growth by free interface diffusion, Proc. Natl. Acad. Sci. 99 (2002) 16531–16536. doi:10.1073/pnas.262485199.

[11] J.S. Marcus, W.F. Anderson, S.R. Quake, Microfluidic single-cell mRNA isolation and analysis, Anal. Chem. 78 (2006) 3084–3089. doi:10.1021/ac0519460.

[12] A.M. Streets, X. Zhang, C. Cao, Y. Pang, X. Wu, L. Xiong, L. Yang, Y. Fu, L. Zhao, F. Tang, Y. Huang, Microfluidic single-cell whole-transcriptome sequencing, Proc. Natl. Acad. Sci. 111 (2014) 7048–7053. doi:10.1073/pnas.1402030111.

[13] M.A. Unger, H.P. Chou, T. Thorsen, A. Scherer, S.R. Quake, Monolithic microfabricated valves and pumps by multilayer soft lithography., Science. 288 (2000) 113–116. doi:10.1126/SCIENCE.288.5463.113.

[14] D.C. Duffy, J.C. McDonald, O.J.A. Schueller, G.M. Whitesides, Rapid prototyping of microfluidic systems in poly(dimethylsiloxane), Anal. Chem. 70 (1998) 4974–4984. doi:10.1021/ac980656z.

[15] S. Satyanarayana, R.N. Karnik, A. Majumdar, Stamp-and-stick room-temperature bonding technique for microdevices, J. Microelectromechanical Syst. 14 (2005) 392–399. doi:10.1109/JMEMS.2004.839334.

[16] S. Thompson, A.R. Abate, Adhesive-based bonding technique for PDMS microfluidic devices3, Lab Chip. 13 (2013) 632–635. doi:10.1039/c2lc40978j.

[17] K. Haubert, T. Drier, D. Beebe, PDMS bonding by means of a portable, low-cost corona system, Lab Chip. 6 (2006) 1548. doi:10.1039/b610567j.

[18] O.C. Jeong, T. Yamamoto, S.W. Lee, T. Fujii, S. Konishi, Surface Modification, Mechanical Property, and Multi-Layer Bonding of PDMS and its Applications, 9th Int. Conf. Miniturized Syst. Chem. Life Sci. 1 (2005) 202–204.

[19] M.A. Eddings, M.A. Johnson, B.K. Gale, Determining the optimal PDMS-PDMS bonding technique for microfluidic devices, J. Micromechanics Microengineering. 18 (2008) 067001. doi:10.1088/0960-1317/18/6/067001.

[20] K. Khanafer, A. Duprey, M. Schlicht, R. Berguer, Effects of strain rate, mixing ratio, and stress–strain definition on the mechanical behavior of the polydimethylsiloxane (PDMS) material as related to its biological applications, Biomed. Microdevices. 11 (2009) 503–508. doi:10.1007/s10544-008-9256-6.

[21] Z. Wang, A.A. Volinsky, N.D. Gallant, Crosslinking effect on polydimethylsiloxane elastic modulus measured by custom-built compression instrument, J. Appl. Polym. Sci. 131 (2014). doi:10.1002/app.41050.

[22] P.M. Fordyce, C.A. Diaz-Botia, J.L. Derisi, R. Gomez-Sjoberg, Systematic characterization of feature dimensions and closing pressures for microfluidic valves produced via photoresist reflow, Lab Chip. 12 (2012) 4287–4295. doi:10.1039/c2lc40414a.

[23] J.N. Lee, X. Jiang, D. Ryan, G.M. Whitesides, Compatibility of mammalian cells on surfaces of poly(dimethylsiloxane), Langmuir. 20 (2004) 11684–11691. doi:10.1021/la048562+.

[24] J.C. McDonald, G.M. Whitesides, Poly(dimethylsiloxane) as a material for fabricating microfluidic devices, Acc. Chem. Res. 35 (2002) 491–499. doi:10.1021/ar010110q.

[25] J.A. White, A.M. Streets, Controller for microfluidic large-scale integration, HardwareX. (2017). doi:10.1016/j.ohx.2017.10.002.

[26] F. Awaja, Autohesion of polymers, Polymer (Guildf). 97 (2016) 387–407. doi:10.1016/J.POLYMER.2016.05.043.

